# Spatial genome re-organization between fetal and adult hematopoietic stem cells

**DOI:** 10.1101/628214

**Authors:** C Chen, W Yu, J Tober, P Gao, B He, K Lee, T Trieu, GA Blobel, NA Speck, K Tan

## Abstract

Fetal hematopoietic stem cells (HSCs) undergo a developmental switch to become adult HSCs. The functional properties of the HSCs change dramatically during this switch, including their cycling behavior, hematopoietic lineage outputs and proliferation rate. The relationship between three-dimensional (3D) genome organization, epigenomic state, and transcriptome is poorly understood during this critical developmental transition. Here we conducted a comprehensive survey of the 3D genome, epigenome and transcriptome of fetal and adult HSCs in mouse. We found that chromosomal compartments and topologically associating domains (TAD) are largely conserved between fetal and adult HSCs. However, there is a global trend of increased compartmentalization and TAD boundary strength in adult HSCs. In contrast, dynamics of intra-TAD chromatin interactions is much higher and more widespread, involving over a thousand gene promoters and distal enhancers. Such dynamic interactions target genes involved in cell cycle, metabolism, and hematopoiesis. These developmental-stage-specific enhancer-promoter interactions appear to be mediated by different sets of transcription factors in fetal and adult HSCs, such as TCF3 and MAFB in fetal HSCs, versus NR4A1 and GATA3 in adult HSCs. Loss-of-function studies of TCF3 confirms the role of TCF3 in mediating condition-specific enhancer-promoter interactions and gene regulation in fetal HSCs. In summary, our data suggest that the fetal-to-adult transition is accompanied by extensive changes in intra-TAD chromatin interactions that target genes underlying the phenotypic differences between fetal and adult HSCs.

## Introduction

During development, hematopoietic stem cells (HSCs) first appear in major arteries of the mouse embryo at embryonic day 11 (E11) and migrate to the fetal liver (FL) at E12 where they expand in number by 10-to 30-fold (*1*). Right before birth, FL HSCs migrate to bone marrow (BM) to take up permanent residence. The physiological properties and functions of FL and adult BM HSCs are distinct. FL HSCs must support rapid blood development and hence rapidly expand, while BM HSCs support homeostatic blood production, and respond to injury and external stress. Phenotypic differences between FL and BM HSCs enable them to fulfill these different physiological needs. Most (>70%) BM HSCs exist in a quiescent G_0_ state (*2, 3*) to prevent HSC exhaustion, whereas the majority of FL HSCs are actively cycling (*4*). Second, the relative lineage outputs of lymphoid and myeloid cells change between FL and BM HSCs and during the process of aging. FL HSCs tend to have balanced lymphoid and myeloid lineage outputs whereas BM HSCs tend to have a myeloid-biased lineage output that becomes more prevalent during the aging process in mouse (*5, 6*). Finally, FL HSCs more robustly engraft mice when transplanted, and display a greater self-renewal activity when stimulated to proliferate *in vivo*. FL HSCs regenerate daughter HSCs in irradiated recipients more quickly and outcompete the production of HSCs from adult BM HSCs (*7, 8*). These phenotypic differences correlate with changes in HSC gene expression (*9*), indicating that FL and BM HSCs are sustained by distinct transcriptional programs.

The three-dimensional genome organization plays an important role in transcription via multiple mechanisms, from long-range interactions between gene promoters and enhancers to higher-order chromosome compartments and domains that can act as expression domains (*10, 11*). Global reorganization of 3D genome structures have been studied in different developmental systems, including human ES cell differentiation(*12*), mouse neural development(*13*), B cell reprogramming process (*14*), T cell linage commitment(*15*), and fetal vs adult erythroid cells (*16*) revealing new insights into the interplay between genome organization, gene expression, and cellular identity.

To date, little is known about how 3D genome organization contributes to the phenotypic difference between fetal and adult HSCs. Better understanding of the 3D genome organization may provide new ways to manipulate gene expression and HSC behavior for translational research. Here, we characterize the phenotypic differences between FL and BM HSCs. We next map the differences in 3D genome organization, epigenomic state, and gene expression between FL and BM HSCs. We reveal a general trend of increased dynamics in 3D genome organization as one moves down the organizational hierarchy. Moreover, our data suggests a high degree of intra-TAD promoter interactome dynamics during the fetal-to-adult HSC transition. We further identify a set of transcription factors potentially involved in mediating developmental-stage-specific promoter-enhancer interactions.

## Results

### Differences in hematopoietic lineage potential and cell cycle status between fetal and adult HSCs

To examine the difference in lineage potentials of fetal and adult HSCs, we purified fetal and adult murine HSCs from embryonic (E) day 14.5 FL and adult (6-8 wks) BM using the cell surface phenotype Lin^-^ Sca-1^+^ c-Kit^+^ CD135^-^ (LSKCD135^-^, Fig. 1A) (*17-21*). We analyzed colonies derived from single HSC for the generation of granulocytes-macrophages (GM), B, and T cells (Fig. 1B, C). Among all plated cells, 33% and 39% cells produced colonies (Fig. 1D). Of the single-cell-derived colonies with lineage readout (Fig. 1D), ∼53% produced mature cells of all three GM, B, and T cells (GM/B/T) for both FL and BM HSCs (Fig. 1E). We found that FL HSCs gave rise to more lymphoid clones (B and/or T cells) whereas adult HSCs gave rise to more GM clones (p < 0.05, Fig. 1E). To determine the long-term multilineage potential of the prospectively purified HSCs, we transplanted purified FL/BM HSCs competitively into lethally irradiated recipient mice. We analyzed GM, B and T cells in peripheral blood using flow cytometry16 weeks post-transplantation. The transplanted FL and BM HSCs engrafted 7/9 and 8/10 of the mice respectively. Moreover, consistent with our single-cell analysis, peripheral blood reconstituted from BM HSCs has a higher GM/(B+T) ratio than that reconstituted from FL HSCs, again suggesting a myeloid lineage bias in BM HSCs (Fig. 1F). Taken together, these results demonstrate the multilineage potential of the HSCs purified in this study and confirm previous reports that BM HSCs have a myeloid lineage bias.

**Fig. 1.**
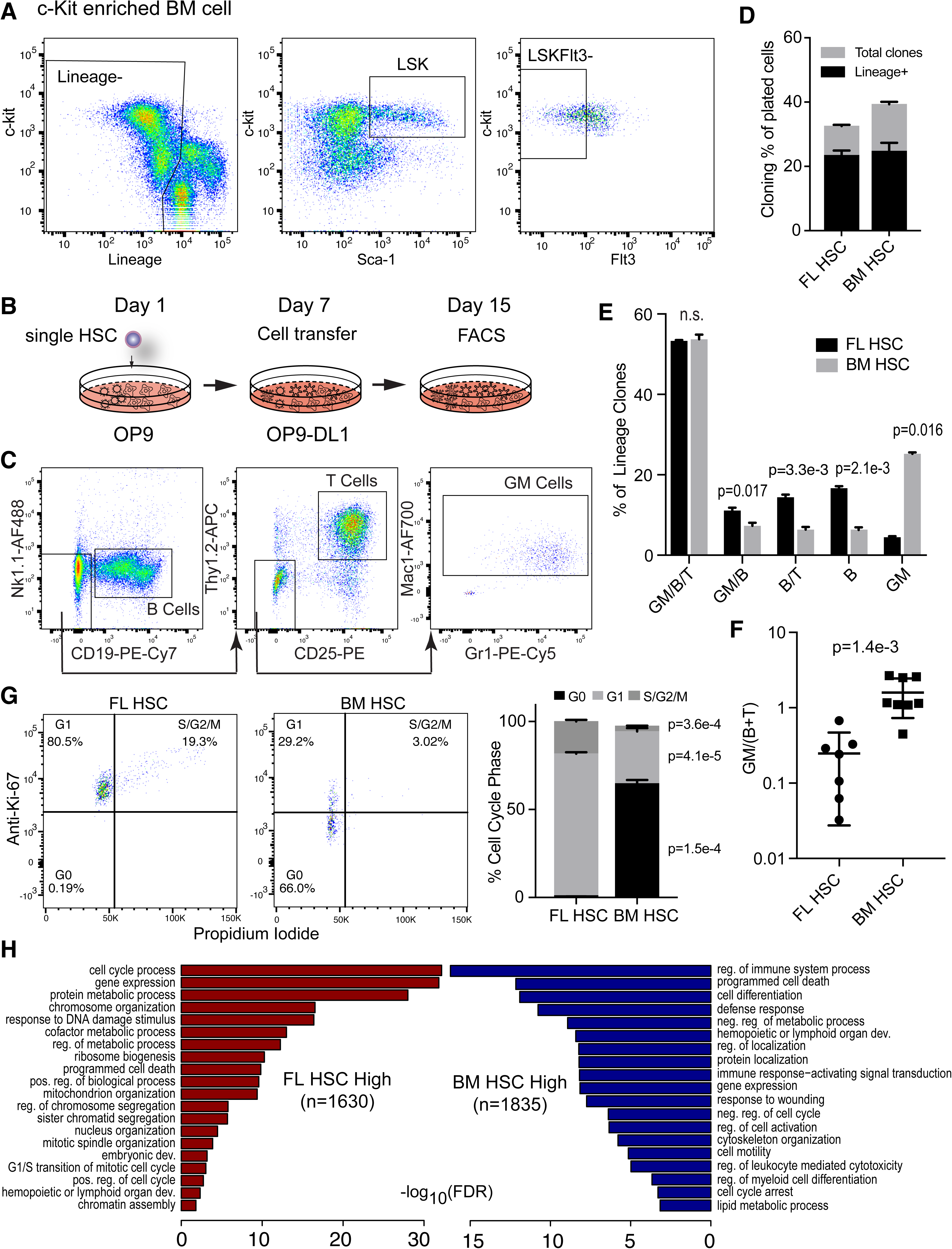
Differences in cell cycle and lineage potential between fetal and adult HSCs. **A)** Representative FACS plots for the purification of adult bone marrow HSCs. **B)** Schematic for the assay for determining combined lineage potentials of HSCs. Single cells are FACS sorted directly into wells with OP9 cells. After 7 days of co-culture, all cells in a clone are transferred to OP9-DL1 cells. Clones are analyzed for lineages after 15 days of co-culture. **C)** Representative FACS plots of a fetal/adult single-HSC-derived GM/B/T clone. A clone is scored positive for B cells if it contains NK1.1^-^CD19^+^ cells; T cells if containing NK1.1^-^CD19^-^CD25^+^Thy1.2^hi^ cells and GM cells if containing NK1.1^-^ CD19^-^CD25^-^Thy1.2^-^Mac1^+^Gr1^-/+^ cells. **D)** Cloning frequency of single HSCs. White bar; total clones; black bar, clones assigned to the GM, B, and/or T lineage. Values are mean ± s.d. of two biological replicates. Each replicate consists of 192 single cells. **E)** Lineage (GM/B/T) composition of lineage-positive clones derived from single fetal or bone marrow HSC cells. **F)** Lineage composition in peripheral blood from transplantation experiment. 50 FL/BM HSCs were competitively transplanted into lethally irradiated wild-type recipients. Long-term multilineage potentials were determined by analyzing peripheral blood at 16 weeks post transplantation. Y-axis, GM/(B+T) ratio in peripheral blood. **G)** Quantification of cell cycle phases by co-staining with anti-Ki-67 antibody and propidium iodide. Left, representative FACS plots. Right, quantification of fractions of HSCs in different cell cycle phases, mean ± s.d. of three biological replicates. P-values were computed using t-test. **H)** Enriched GO terms among genes up-regulated in either FL HSCs or BM HSCs. Numbers of differentially expressed genes are indicated in the parenthesis.

To examine the cell cycle difference between FL and BM HSCs, we co-stained FL and BM samples with HSC surface markers (LSKCD135^-^), anti-Ki-67 antibody and propidium iodide (Fig. 1G), to simultaneously purify HSCs and profile their cell cycle status. Consistent with previous studies, the fractions of cells in each cell cycle phase are significantly different between FL and BM HSCs. In particular, the vast major of FL HSCs are cycling (only 0.19% in G_0_ phase) whereas the majority of BM HSCs (66%) are in G_0_ phase (Fig. 1G).

Next, we performed RNA-Seq to profile the transcriptomes of fetal and adult HSCs. Using a False Discovery Rate cutoff of 0.05 and fold change cutoff of 2, we identified 3,464 differentially expressed genes, including 1630 and 1834 genes expressed higher in either FL HSCs or BM HSCs, respectively (Supplemental Table S1). Genes expressed higher in FL HSCs are enriched for Gene Ontology terms of “cell cycle process”, “ribosome biogenesis”, “G1/S transition of mitotic cell cycle”, “chromosome organization”, consistent with the larger fraction of FL HSCs in cell cycle (Fig. 1H). In contrast, genes expressed higher in BM HSCs are enriched for GO terms of “regulation of immune response”, “cell differentiation”, “cell cycle arrest”, “hematopoietic or lymphoid organ development”, consistent with the phenotypic differences.

### Chromosome compartments are largely unchanged, but compartmentalization is strengthened during the fetal-to-adult transition

To compare the global 3D genome organizations, we profiled genome-wide chromatin interactions using *in situ* Hi-C (*22*). We also conducted Capture-C (*23*) to investigate the dynamics of promoter-centric chromatin interactions, focusing on 4,052 promoters that are highly expressed in HSCs compared to a compendium of 20 other mouse tissues (Fig. 2A, Methods, Supplemental Fig. S1, Supplemental Table S2). Using several metrics, we confirmed that our Hi-C and Capture-C data have sufficient sequencing depth and high reproducibility (Supplemental Fig. S2, Supplemental Table S3). To understand the relationship between the epigenome and 3D genome organization, we also generated ChIP-Seq data for four histone marks, H3K4me1, H3K4me3, H3K27ac, and H3K27me, as well as ATAC-Seq data (Fig. 2A). These data also have sufficient sequencing depth and high reproducibility (Supplemental Fig. S3,S4).

**Fig. 2.**
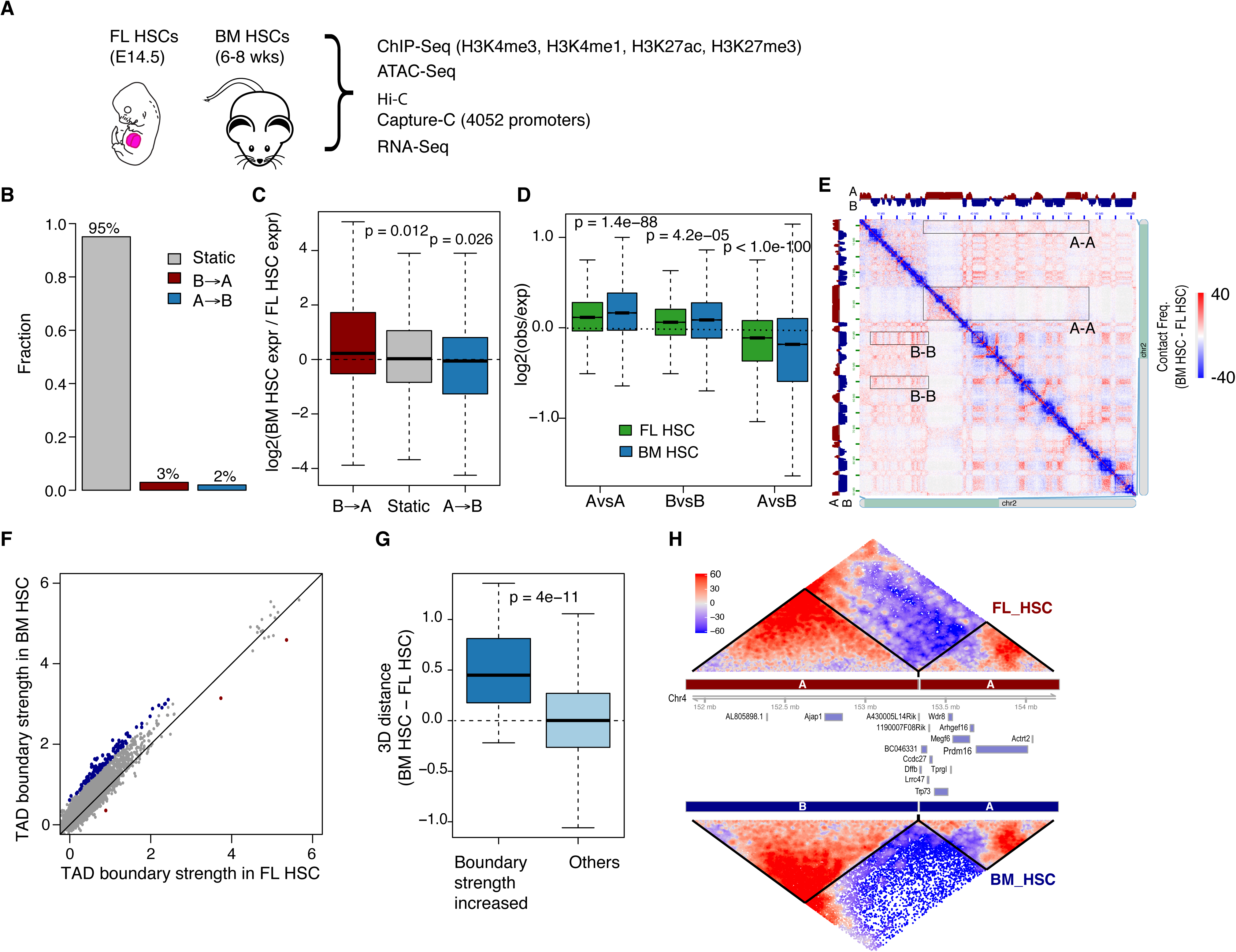
Limited change in global 3D genome organization during fetal-to-adult HSC transition. **A**) Schematic diagram of experimental design. **B)** Fraction of genomic regions with compartment switching during fetal to adult transition. B→A, regions switching from compartment B to compartment A; static, regions without compartment switching. **C**) Gene expression change is correlated with compartment switching. **D-E**) Increased compartmentalization during fetal to adult transition. **D)** Shown are log ratios of observed versus expected contact frequencies between TADs from the same (A vs A, B vs B) or different compartments (A vs B). **E)** An example heatmap of contact frequencies along chromosome 2, showing increased contacts among regions of the same compartment. Compartment assignment is indicated along the top and left. Several examples of more frequent interactions between the same compartments are highlighted by rectangles. Color is proportional to the difference in contact frequency (BM HSC - FL HSC). **F)** Scatter plot of TAD boundary strength in FL HSCs and BM HSCs. Boundaries with significantly increased and decreased strength (FDR < 0.1) are highlighted in blue and red, respectively. **G)** 3D distance is larger between adjacent TADs with increased boundary strength during the fetal-to-adult transition. Y-axis, difference in 3D distance of adjacent TADs between BM HSCs and FL HSCs. Normalized distance was calculated based on the 3D structure model of each chromosome. **H)** An example TAD boundary with significantly increased strength during the transition. TAD heatmap color is proportional to SHAMAN score. P-values in panels C), D), G) were calculated using t-test.

Previous studies have revealed that the 3D genome is organized in a hierarchical fashion. At the top level are so-called A and B compartments with an average size of 3 Mb. The A compartments correlate with early replicating, euchromatic regions whereas the B compartments correlate with heterochromatin (*12, 24*). We found that in both FL and BM HSCs, the genome is equally divided into the A and B compartments (Supplemental Fig. S5A, S5B). Compartment A is associated with higher levels of active epigenetic marks including H3K4me3, H3K4me1 and H3K27ac, higher chromatin accessibility, and higher gene expression. In contrast, compartment B is associated with lower chromatin accessibility and lower gene expression (Supplemental Fig. S5C, S5D). Overall, 95% of the genome falls into the same compartment in both FL and BM HSCs, suggesting limited change in the global 3D genome organization during the fetal-to-adult HSC transition (Fig. 2B). Of the 5% of the genome that switches compartment, genes are expressed significantly higher when the corresponding region switches from compartment B to compartment A and vice versa (Fig. 2C). Despite the limited change in the locations of compartment boundaries, the overall compartmentalization is strengthened in BM HSCs, as indicated by the increased interactions among TADs from the same compartments and decreased interactions among TADs from different compartments in BM HSCs (Fig. 2D, E).

### Locations of TADs boundaries are largely conserved but boundary strength increases during fetal-to-adult transition

Topologically associating domains (TADs) ^17,18^, ranging in size from 100kb to 3Mb and average 1Mb, have been suggested to be important organization units of the 3D genome. Using the GMAP algorithm (*25*), we identified 2,393 and 2,391 TADs in FL and BM HSCs, respectively. Similar to compartment boundaries, 88% of TAD boundaries are shared between the two cell types (Supplemental Fig. S6A). Although the change in boundary location is small, we observed a global trend of increased TAD boundary strength, as measured by the difference in intra-TAD and inter-TAD interactions (see Methods for detail), during the fetal-to-adult transition (Supplemental Fig. S6B). This observation is further supported by our Capture-C data since the fraction of inter-TAD promoter-centric interactions is significantly reduced during the transition (Supplemental Fig. S6C). This global increase in boundary strength could be due to the difference in the fraction of cells in different cell cycle phases. A recent single-cell Hi-C study suggests that TAD boundary strength is highest in G_1_ phase and decreases as the cell enters S phase (*26*). By comparing published Hi-C data of pure proliferating and quiescent human fibroblast cells, we also found that TAD boundary strength is higher in the quiescent cells than in proliferating cells (Supplemental Fig. S7A) (*27*). Taken together, these data suggest that the higher fraction of G_0_ cells (Fig. 1G) in BM HSCs potentially account for the higher TAD boundary strength in these cells. Besides this general trend of increased boundary strength, we found 58 (3%) TADs whose boundaries exhibit significantly increased strength in BM HSCs compared to FL HSCs (FDR < 0.1, Fig. 2F).

Interestingly, 3D modeling of the genome shows that the two adjacent TADs of those boundaries are farther apart in BM HSCs than in FL HSCs (Methods, Fig. 2G), suggesting a positive correlation between boundary strength and physical distance between adjacent TADs. An example TAD boundary with significant increased strength is shown in Fig. 2H. Additional examples are shown in Supplemental Fig. S8A-H. Using our Capture-C data, we also found inter-TAD enhancer-promoter interactions across those boundaries are significantly reduced (Supplemental Fig. S9A, S9B), suggesting the stronger boundaries may further impede the inter-TAD enhancer-promoter interactions in BM HSCs.

### Intra-TAD promoter interactome exhibits substantial dynamics during fetal-to-adult transition

The analyses above suggest limited change of the 3D genome at the compartment and TAD levels. We therefore investigated the dynamics of chromatin interactions within TADs. We used our Hi-C data to identify TADs and Capture-C data to identify promoter-centric interactions. To identify statistically significant Capture-C interactions, we developed the LiMACC algorithm (Local iterative Modeling Approach for Capture-C data) (Methods). Performance benchmarking shows that LiMACC has better performance compared to the state-of-the-art method, CHiCAGO(*28*) (Supplemental Fig. S10), in terms of identifying higher fractions of functional interactions, including enhancer-promoter interactions, promoter-promoter interactions and promoter-ATAC-Seq peak interactions. At an FDR cutoff of 0.01, 89,545 significant interactions were identified (Supplemental Table S5). Among them, 32,814 (36.6%) are FL HSC-specific, 29,209 (32.6%) are BM HSC-Specific, and 27,522 (30.8%) are shared (Supplemental Fig. S11).

By comparing the interaction frequencies of the set of promoters in the same TAD, we identified 242 TADs exhibiting significant dynamics of the intra-TAD promoter interactome between FL and BM HSCs (Fig. 3A, Supplemental Table S4, Methods). Genes in these TADs are enriched for GO terms such as cell cycle, metabolism, chromosome maintenance and apoptosis (Fig. 3B), consistent with the phenotypic difference between FL and BM HSCs. Interestingly, we found dynamic TADs are marked by significantly higher levels of active chromatin marks (H3K4me1, H3K4me3 and H3K27ac) and higher chromatin accessibility, compared to static TADs (Supplemental Fig. S12). A dramatic example of such dynamic TADs is located around the *Hmga2* locus. *Hmga*2 has multiple roles including chromatin architectural protein and transcription factor. It has been shown to play a critical role in the higher level of self-renewal activity in FL HSCs (*29*). We found that multiple interactions involving the *Hmga2* promoter and two super enhancers in FL HSCs disappear in BM HSCs. This loss of enhancer interaction is associated with a significant decrease in *Hmga2* expression (Fig. 3C, 3E) in BM HSCs. Previous studies have shown that *Hmga2* is post-transcriptionally regulated by the *Lin28b*-*let7* axis (*29*). Our data suggests *Hmga2* is also regulated at the chromatin interaction and transcriptional level. The other gene *Llph* in this TAD also loses three enhancer interactions but interestingly gains two promoter interactions in BM HSCs. Another example of dynamic intra-TAD interaction involves the *Smarca2* gene, a member of the SWI/SNF chromatin remodeling complex (Fig. 3D). The *Smarca2* promoters gain several interactions with active enhancers and *Smarca2* expression is up-regulated during the fetal-to-adult transition (Fig. 3E). Additional examples are shown in Supplemental Fig. S13 and S14. Taken together, these data suggest that intra-TAD chromatin interaction dynamics plays a major role in driving the phenotypic differences between FL and BM HSCs.

**Fig. 3.**
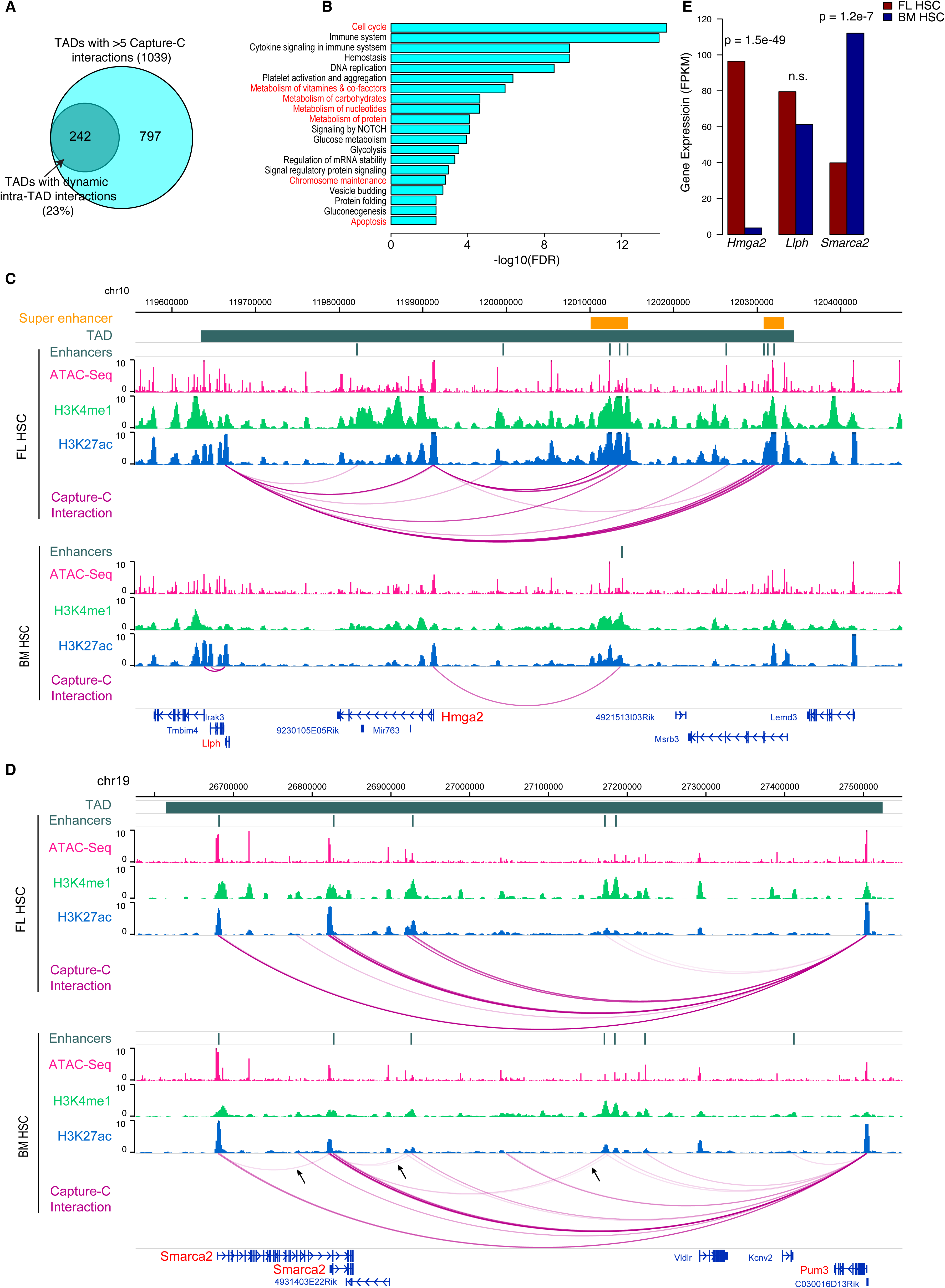
Intra-TAD promoter-centric interactions exhibit large dynamics. **A)** Venn diagram of TADs with dynamic intra-TAD interactions during fetal-to-adult HSC transition. **B)** Enriched GO terms among genes in the TADs with dynamic intra-TAD interactions. **C)** An example TAD with more promoter-centric interactions in FL HSCs than BM HSCs. Gene promoters with Capture-C baits are highlighted in red. TAD is indicated with a navy green bar. The normalized signals of ATAC-Seq, H3K4me1, H3K27ac, and Capture-C are displayed for FL HSCs (upper tracks) and BM HSCs (lower tracks). Two super enhancers are indicated with an orange bar. **D)** An example TAD with more promoter-centric interactions in BM HSCs than FL HSCs (indicated by arrows). **E)** Expression levels of three genes, *Hmga2, Llph* and *Smarca2* in the TAD. P-value for differential expression was computed using the edgeR software.

### Dynamic enhancer-promoter interactions target genes underlying phenotypic differences

Using our histone mark ChIP-Seq data and the CSI-ANN algorithm (*30*), we identified active enhancers and promoters in both cell types (Methods and Supplemental Fig. S15). About 20% and 30% of the 89,545 significant promoter-centric interactions are Enhancer-Promoter (EP) and Promoter-Promoter (PP) interactions (Supplemental Fig. S11B), respectively. Among the EP interactions, 57% are cell-type-specific (Fig. 4A). Genes targeted by cell-type-specific EP interactions are expressed significantly higher in the same cell type (p=7.8e-6, Fig. 4B). They are also involved in multiple biological processes that underlying the phenotypic differences (Fig. 4C).

**Fig. 4.**
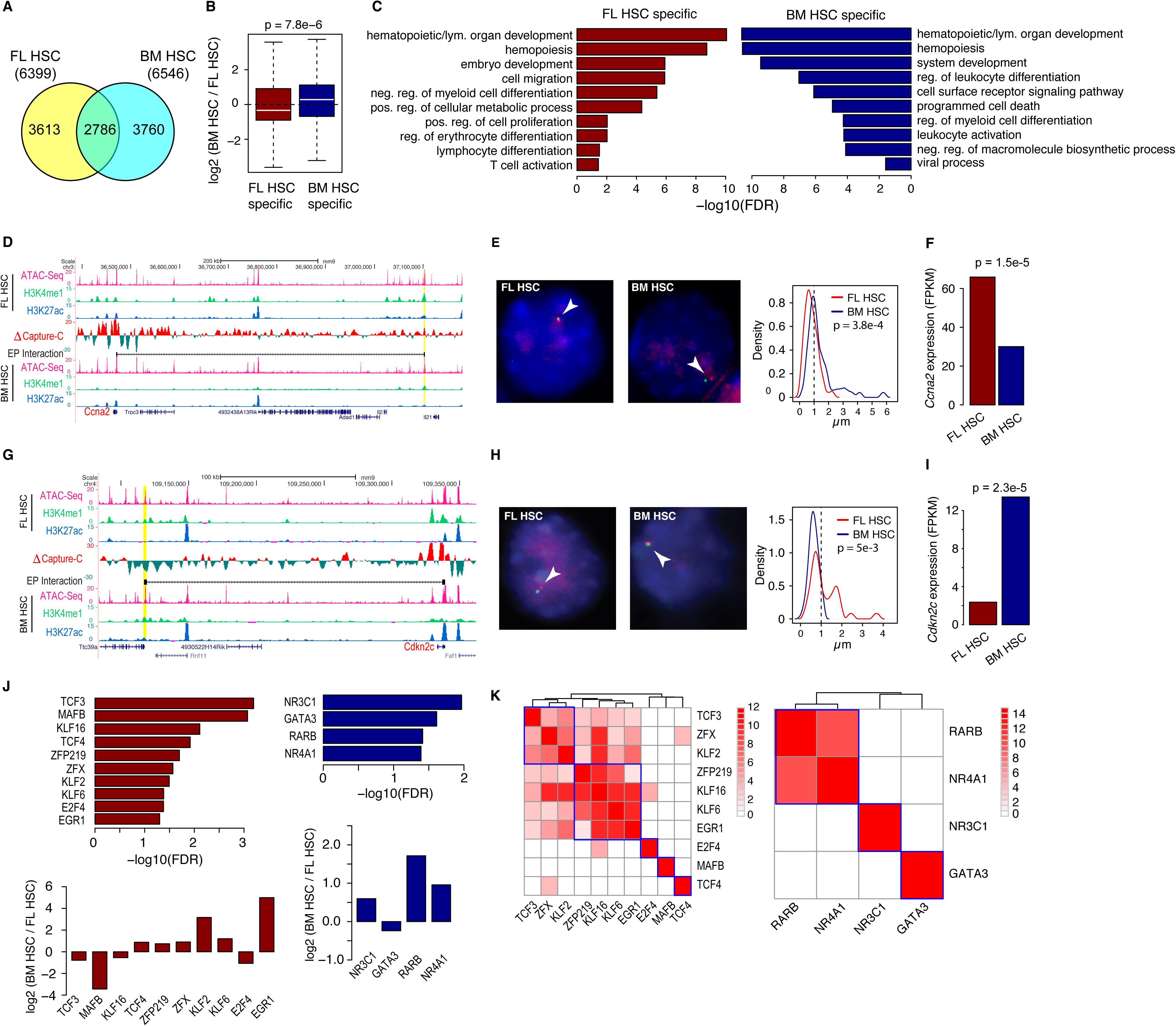
Dynamic Enhancer-Promoter (EP) interactions account for phenotypic differences between FL and BM HSCs. **A)** Venn diagram of EP interactions detected by Capture-C. >60% EP interactions are cell type-specific. **B)** Expression change of genes with cell-specific EP interactions. Expression change was calculated as FPKM ratio of BM HSCs to FL HSCs. P value was calculated using t-test. **C)** Enriched GO terms of genes with FL HSC-specific and BM HSC-specific EP interactions. **D)** An example of FL HSC-specific EP interactions involving the promoter of *Ccna2*. Difference in normalized Capture-C signal is shown in the middle track. Normalized ATAC-Seq signal, H3K4me1 and H3K27ac ChIP-Seq signals are displayed in the rest of the tracks. Gene whose promoter was used as Capture-C bait is marked as red. **E)** DNA FISH confirms the *de novo* FL HSC-specific EP interaction. Left, representative DNA FISH images of the *Ccna2* promoter (red) and enhancer (green) in FL HSCs (left panel) and BM HSCs (right panel). Interaction is denoted by a white arrow. Right, frequency of the quantified distance distribution between *Ccna2* promoter and the enhancer *μ*m) (# nuclei imaged: 90 and 59 for FL HSC and BM HSC, respectively). P value was calculated using t-test. **F)** Gene expression level of *Ccna2*. P-value of differential expression was calculated using edgeR. **G)** An example of BM HSC-specific EP interaction involving the promoter of *Cdkn2c*. **H)** DNA FISH confirmation of the EP interaction. **I)** Gene expression level of *Cdkn2c*. **J)** Enriched TF DNA binding motifs at enhancers of FL HSC-specific and BM HSC-specific EP interactions. Bottom plots, expression levels of the TFs with enriched motifs. **K)** Co-localization of enriched TF motifs at enhancers of cell-specific EP interactions. Color of heatmap is proportional to the p-value of co-localization. Heatmap was clustered using hierarchical clustering.

An example FL HSC-specific EP interaction involving the gene *Ccna2* is shown in Fig. 4D-4F. *Ccna2* gene is a positive regulator of G_1_/S and G_2_/M transitions and is expressed significantly higher in FL HSCs (Fig. 4F). An active enhancer located 700Kb downstream of the *Ccna2* promoter forms an interaction with the *Ccna2* promoter in FL HSCs but not in BM HSCs. An example of BM HSC-specific EP interaction involving the gene *Cdkn2c* is shown in Fig. 4G-4I. *Cdkn2c* is a negative regulator of cell cycle G_1_ phase progression. DNA fluorescence *in situ* hybridization (DNA-FISH) confirms both cell-specific EP interactions (Fig. 4E and 4H). Additional examples are shown in Supplemental Fig. S16.

To identify transcription factors that are involved in cell-type-specific EP interactions, we conducted TF motif analysis of enhancers involved in cell-specific EP interactions. We identified 22 and 6 TFs whose motifs are enriched at enhancers of FL HSC-specific and BM HSC-specific EP interactions (p < 0.01), respectively. Among those TFs, 10 and 4 are differentially expressed in FL and BM HSCs, respectively (Fig. 4J). Moreover, many of the enriched TFs have co-localized binding sites within the same enhancers involved in the EP interactions (Fig. 4K), suggesting combinatorial binding of these TFs may be required for the stage-specific EP interactions.

### TCF3 mediates developmental-stage-specific enhancer-promoter interactions

Transcription factor 3 (TCF3, also known as E2A) ranks as the top TF whose motif is enriched at enhancers of FL HSC-specific EP loops (Fig. 4J). TCF3 is required for B and T cell development (*31*) and HSC maintenance (*32*). It is also implicated in chromatin organization in B cells (*33, 34*). We found *Tcf3* is expressed 1.7-fold higher in FL compared to BM HSCs (p=7e-3). FL HSC-specific TCF3 targets (genes targeted by FL HSC-specific EP loops that have TCF3 DNA binding sites in the enhancers) were also expressed higher in FL HSCs (Fig. 5A). We confirmed TCF3 binding to a number of these enhancers using ChIP-qPCR (Fig. 5B). The FL HSC-specific TCF3 targets are enriched for GO terms such as ‘cell cycle phase’, ‘chromatin organization’, ‘regulation of cell proliferation’, and ‘lymphocyte activation’. Taken together, these data suggest that TCF3 occupies FL HSC specific EP loops and regulates the expression of genes underlying the phenotypic differences.

**Fig. 5.**
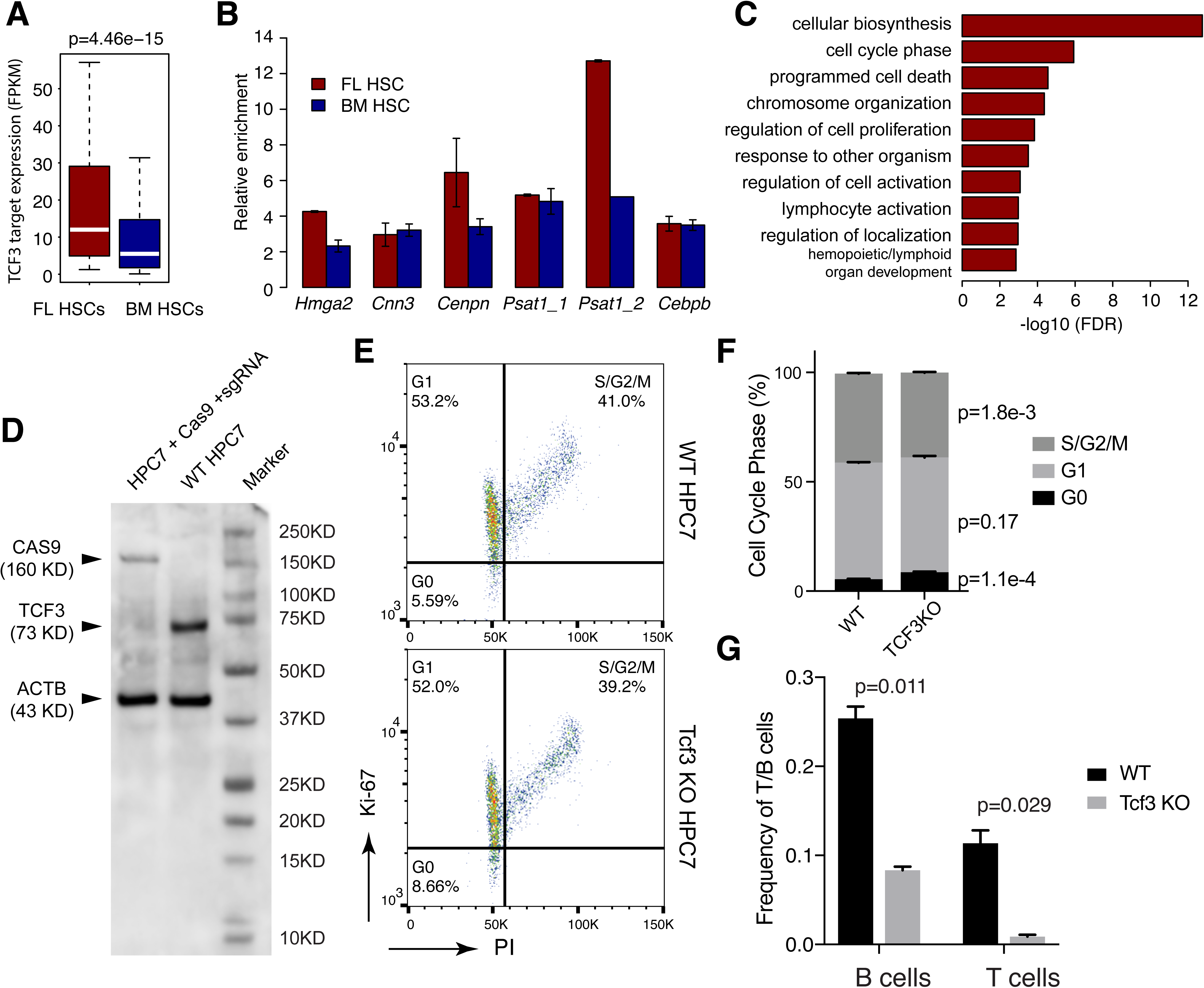
TCF3 occupies developmental-stage-specific enhancer-promoter loops and affects cell cycle phase and lineage potential. **A)** FL HSC-specific TCF3 targets have significantly higher expression. **B)** ChIP-qPCR confirmation of TCF3 binding to enhancers involved in FL HSC-specific EP loops. **C)** Enriched GO terms among genes targeted by FL HSC-specific EP loops occupied by TCF3. **D)** Western blot showing knock down of TCF3 by CRISPR-Cas9**. E-F)** Cell cycle phase analysis by co-staining with propidium iodide and anti-Ki-67 antibody. **E**) representative FACS plots of wild type and Tcf3 knockout HPC-7 cells. **F**) quantification of cell cycle phases, mean ± s.d. of three biological replicates. P-values were computed using t-test. **G)** Limiting dilution assay showing significantly reduced lymphoid potential in Tcf3 knockout HPC-7 cells. Y-axis, frequencies of CD45^+^CD25^+^CD90^+^ T progenitors and CD45^+^B220^+^CD19^+^ B progenitors produced by wild type and Tcf3 knockout HPC-7 cells after 10∼12 days of co-culturing with OP9/OP9-DL1 cells. Values are mean of 2 biological replicate experiments. Error bar, standard deviation.

To further confirm if TCF3 can mediate developmental-stage-specific enhancer-promoter interactions, we performed Capture-C experiment comparing wild type HPC-7 cells and HPC-7 cells with *Tcf3* knockout using CRISPR-Cas9 technology (Fig. 5D). HPC-7 is a murine multi-potent hematopoietic precursor cell line that has been used as a model of HSCs (*35-37*). *Tcf3* knockout has a modest effect on the overall cell cycle duration of HPC-7 cells (Supplemental Fig. S17). However, it significantly increases the fraction of cells in G_0_ phase (p = 1.1e-4) and reduces the fraction of cells in G_2_/S phase (p = 1.8e-3) (Fig. 5E). Using limiting dilution assay, we further investigated the differentiation potential of HPC-7 cells with *Tcf3* knockout. We found that the lymphoid potential of these cells is dramatically reduced (Fig. 5G). The frequencies of CD45^+^CD19^+^B220^+^ B cells and CD45^+^CD25^+^CD90^+^ T cells are reduced from 1/4 to 1/12 (p = 0.011) and from 1/9 to 1/115 (p = 0.029), respectively. Taken together, these results suggest that TCF3 plays a role in cell cycle and lymphoid potential in HPC-7 cells.

We found that knocking out *Tcf3* significantly reduces the contact frequency of enhancer-promoter interactions in which the enhancers are occupied by TCF3 (n=93, Fig. 6A) in FL HSCs. Of these FL HSC-specific EP loops that are bound by TCF3, several of them target key TFs in FL HSCs, such as *Hmga2*; cell cycle genes, such as *Rcc2* (regulator of chromosome condensation 2), *Cenpn* (centromere protein N); and metabolic genes, such as *Cryl1* (crystallin lambda 1), *Psat1* (phosphoserine aminotransferase 1), *Prkag1* (protein kinase AMP-activated non-catalytic subunit gamma 1) and *Scd1* (Stearoyl-CoA desaturase). Genome browser view of Capture-C signal and TCF3 ChIP-Seq signal for the *Hmga2, Rcc2*, and *Psat1* loci are shown in Fig. 6C-E. Additional examples are provided in Supplemental Fig. S18. The relative expression levels of those genes are significantly decreased in *Tcf3* knockout cells (Fig. 6B, p < 0.05). In summary, these results confirm that TCF3 can mediate developmental-stage-specific enhancer-promoter interactions in fetal HSCs and the genes targeted by these interactions are responsible for fetal HSC specific phenotypes.

**Fig. 6.**
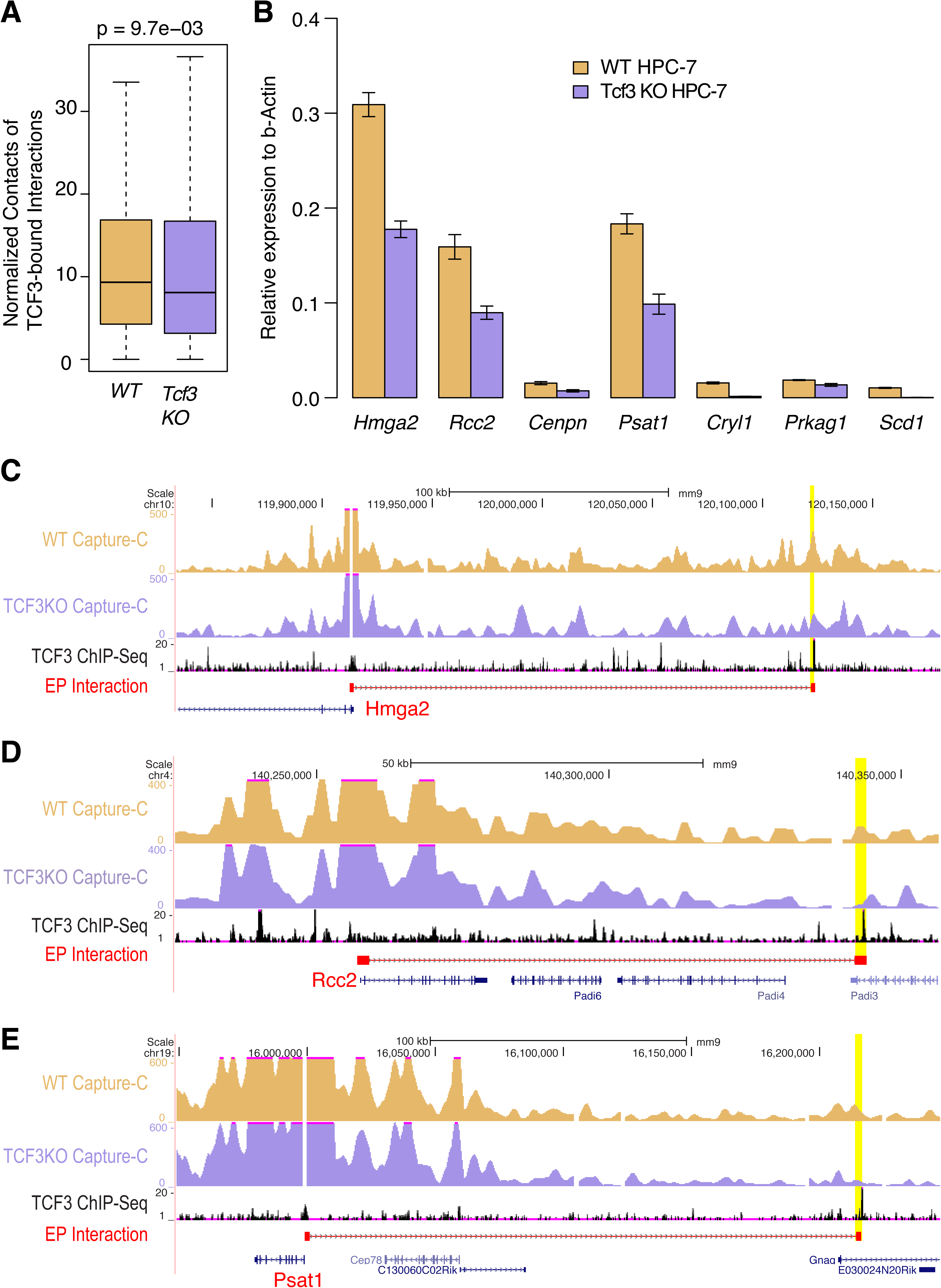
Loss of TCF3 results in loss of cell-specific enhancer-promoter loops and deregulation of target gene expression. **A)** Reduced interaction frequency among TCF3 bound enhancer-promoter loops after knocking down Tcf3. **B)** RT-qPCR of target genes of TCF3 bound enhancer-promoter loops. **C-E)** Capture-C data showing loss of enhancer-promoter interaction after Tcf3 knock down for *Hmga2* (**C**), *Rcc2* (**D**), and *Psat1*(**E**).

## Discussion

Fetal and adult HSCs have dramatic phenotypic differences, especially in their cycling behavior, lineage output and metabolic state. Our RNA-Seq analysis revealed over 3,000 genes that are differentially expressed between these two types of HSCs. Here, we investigated how changes in the different hierarchical levels of 3D genome organization contribute to the differences in gene expression and phenotype. We found an increasing amount of changes going down the genome architectural hierarchy; 5% at the chromosome compartment level, 12% at TADs, 23% at subTADs, and 57% at enhancer-promoter interactions.

Although the location of compartment and TAD remain relatively unchanged during the fetal-to-adult transition of HSCs, we observed a general trend of increased compartmentalization and TAD boundary strength. To further corroborate our finding, we analyzed Hi-C data from four other developmental systems and observed a similar trend (Supplemental Fig. S7), including human fibroblast senescence (*27*), embryonic stem cell (ESC) differentiation to neurons (*13*), differentiation of cardiovascular muscle cells from induced pluripotent stem cells (iPSCs)(*38*) and reprogramming of pre-B cells to pluripotent stem cells (*39*). The mechanism for the increased strength of TAD boundary and compartmentalization during development is unclear. A potential factor may be the cohesin complex. Recent studies have suggested critical and distinct roles of the cohesin complex in compartmentalization versus formation of TADs (*40-42*). Further investigation of different cohesin subunits and other architecture proteins such as condensin (*43*) and their interplay may uncover the mechanisms for the dynamics of compartmentalization and TAD boundaries.

The role of transcription factors in enhancer-promoter interactions is poorly understood during HSC development. Our analysis identified several TFs that are potential mediators of EP interactions in HSCs. We tested one of the predictions, TCF3, by a loss-of-function approach. We observed a significant decrease of FL HSC-specific EP contact frequency in TCF3 knockout HPC-7 cells compared to the wild type HPC-7 cells, suggesting TCF3 is involved in mediating stage-specific EP interactions. Moreover, we identified TCF3 as a novel regulator of *Hmga2*, a key gene distinguishing fetal and adult HSCs. Knockout of *Tcf3* significantly reduces the interactions between *Hmga2* promoter and its enhancers, along with significant downregulation of *Hmga2*, which may contribute to the loss of lymphoid potential of HPC-7 cells.

The observed changes in EP interaction and gene expression after *Tcf3* knockout are significant but modest, suggesting additional TFs might also contribute to EP interactions and phenotypic differences. This is corroborated by the enrichment of colocalized TFs at HSC-specific EP interactions. Multiple lines of evidence further support a potential role in EP interactions for these predicted TFs. For instance, MAFB is known to restrict myeloid lineage choice in HSCs (*44*). We found *Mafb* is expressed 11-fold higher in fetal compared to adult HSCs and it binds to an FL HSC-specific enhancer that targets to the *Igf2* promoter (Supplemental Fig. S19), an important gene that controls FL HSC cycling activity (*45*). Several TFs in the SP/KLF family are also enriched at FL HSC-specific EP interactions. These factors have potential of mediating cell specific EP interactions, for example KLF1 in erythrocytes (*46, 47*) and KLF4 in ESCs (*48*).

EGR1 negatively regulates HSC proliferation and mobilization (*49*). Consistent with this role, *Egr1* expression is >30 fold higher in BM HSCs than FL HSCs. Two nuclear receptors, NR3C1 and NR4A1, are enriched at adult HSC-specific EP interactions. NR4A1 was shown to regulate HSC quiescence in adult HSCs (*50*) (*51*), which is consistent with our and previous findings. NR4A1 was shown to restrict the HSC proliferation through inflammatory response. GATA3 has been shown to promote cell cycle entry and proliferation in murine bone marrow HSCs (*52*).

In summary, our study suggests that the fetal-to-adult transition of HSCs is accompanied by a large-scale promoter interactome change within TADs, impacting many gene pathways relevant to phenotypic differences between the two types of HSCs. The newly identified transcription factors and their target genes via EP interactions may present novel targets for developing protocols for HSC mobilization for therapeutic purposes.

## Materials and Methods

### Mouse strains

Female B6129SF1/J mice were mated with male C57BL/6J mice for FL HSCs. Fetal livers were dissected from embryonic (E) day 14.5 (E14.5) embryos. Pairs of female B6129SF1/J and male C57BL/6J (6-8 weeks old) were dissected for bone marrow HSCs. The Children’s Hospital of Philadelphia Office of the Institutional Animal Care and Use Committee review board approved these studies.

### Purification of HSCs from adult bone marrow and E14.5 fetal liver

Marrow of long bones (tibias and femurs) were flushed out with staining buffer (1×PBS + 2% FBS) and stained with anti-mouse CD16/32 antibodies to block non-antigen-specific binding. Stained cells were washed twice with staining buffer and applied to autoMACS to enrich CD117^+^ cells. Enriched cells were stained with the antibody cocktail against CD117 (c-kit), Ly-6A/E (Sca-1), CD135, Ly6G/Ly-6C (Gr-1), CD11b, TER-119, CD4, CD8a, CD45R/B220, CD3ε and CD11c (Supplemental Table S7) at 4°C for 15 min in the dark. Cells were first subjected to yield sort for live HSC (Lin^-^Sca-1^+^c-Kit^+^CD135^-^)(*19, 21, 53*) and collected into 500 µL 1×IMDM + 20% FBS in a 12×75-mm polystyrene tube. Collected cells were sorted by purity sort using the same gating strategy and sorted into 0.8 ml 1×IMDM+50% FBS in a 1.5 ml DNA LoBind tube. Purity of sorted cells is more than 95%.

Fetal livers were dissected from E14.5 embryos. Single cell suspension was prepared by dissociating mechanically followed by red blood cell lysis. Cells were stained with an antibody cocktail of CD117 (c-kit), Ly-6A/E (Sca-1), CD135, Ly6G/Ly-6C (Gr-1), TER-119, CD4, CD8a, CD45R/B220, CD3ε and CD19 (Supplemental Table S7). Yield sort was followed by purity sort for FL HSCs (Lin^-^Sca-1^+^c-Kit^+^CD135^-^) as described above for BM HSCs.

### Lymphoid limiting dilution assays

Lymphoid limiting dilution assay was performed as previously described with a few modifications (*54*). OP9 or OP-DL1 cells were seeded 1 day before co-culture at 4,000 cells/well in a flat 96-well tissue culture plate. For T cell differentiation, cells were cultured with OP9-DL1 in 1ng/mL IL7 and 5ng/mL Flt3L with serial dilution of 50, 25, 12, 6, 3, 2, 1, 0.5 cells per well. Five replicates per dilution were performed. For B cell differentiation, cells were cultured with OP9 in 10ng/mL IL-7 and 5ng/mL Flt3L with the same serial dilution. At days10-12, cells were stained with CD45, CD19, B220 for B differentiation and CD45, CD25, CD90 for T differentiation. Antibody and Cytokine information is listed in Supplemental Table S7.

### Combined lineage potential assay

Single cells (E14.5 fetal liver HSC and adult bone marrow HSC) were directly sorted onto OP9 stromal cells with 25ng/mL SCF, 25ng/ml FLT3L, and 20ng/mL IL-7. Cultures were transferred from OP9 to OP9-DL1 stromal cells after 7 days of culture with 1ng/mL IL-7 and 5ng/mL FLT3L and analyzed by FACS after a total of 15 days. Clones were defined based on the following markers; B cells, NK1.1^-^ CD19^+^; T cells, NK1.1^-^ CD19^-^ CD25^+^ Thy1.2^hi^; GM cells, NK1.1^-^ CD19^-^ CD25^-^ Thy1.2^-^ Gr1^+/-^Mac1^+^ and NK cells, NK1.1^+^ CD19^-^ CD25^-^ Thy1.2^-^. Antibody and Cytokine information is listed in Supplemental Table S7.

### Transplantation experiment

50 FACS-sorted E14.5 fetal liver HSCs or adult bone marrow HSCs (6-8 wks) were transplanted together with 250,000 competitor CD45.1/CD45.2 spleen (from B6.SJL mice) cells into B6.SJL Ptprca Pep3b/BoyJ mice by retroorbital injection. Host mice were irradiated with two split doses administered 3-4 hours apart of 4.3-4.5 Gy from a Cs-137 source. Peripheral blood (PB) analyses were conducted at 15–17 weeks. Mice were considered reconstituted if ≥ 0.1% donor contribution to total CD45^+^ cells were achieved. Antibody information is listed in Supplemental Table S7.

### Cell Cycle Assay

E14.5 fetal liver cells and bone marrow cells were enriched with anti-CD117 microbeads and labeled with antibody cocktail for HSCs. Cells were then fixed with 4% paraformaldehyde (43368, Alfa Aesar) in PBS, permeabilized with 1% saponin (47036-50G-F, Sigma). Cells were washed with staining buffer and then stained with anti-Ki-67 antibody conjugated with AlexaFluor700, followed by resuspension with 50ug/mL PI solution (421301, Biolegend). Stained cells were analyzed with BD LSRFortessa.

### BrdU incorporation assay

BrdU incorporation assay was performed based on the instruction of FITC BrdU flow kit (559619, BD Pharmingen). Briefly, cells were seeded into 6-well plates 1 day before BrdU incorporation at the density of 5 × 10^5^/*mL*. BrdU (final concentration 10*μM*) was directly added into culture medium. Cells were fixed, permeabilized, DNase treated, and analyzed at 1 hr, 2 hr, 4 hr, 8 hr, 12 hr, and 24 hr.

### *In situ* Hi-C

*In situ* Hi-C was performed based on previous publication with a few modifications(*22*). Half million sorted cells were cross-linked with 1% formaldehyde for 10 min at RT and quenched with glycine. Nuclei were then permeabilized with 250 μL of cold lysis buffer (10 mM Tris, 10 mM NaCl, 0.2% Igepal CA630) and 50 μL of protease inhibitors (P8340, Sigma). Chromatin was digested with 100 units of MboI overnight at 37°C with rotation. Restriction fragment ends were labelled with biotinylated nucleotides and proximity ligation in a small volume with 5 μL of 400 U/μL T4 DNA ligase. After reversal of cross-link, DNA was sheared to a length of 300∼500 bp and size-selected using AMPure XP beads. The ligated junctions were pulled down with 150 μL of Dynabeads MyOne Streptavidin C1 magnetic beads. End repair, A-tailing, and addition of Illumina index adaptors were performed on beads. Libraries were size-selected and purified using AMPure XP beads. Libraries were sequenced with 75bp paired-end reads on an Illumina NextSeq and/or Hiseq 2000.

### Selection of promoters and design of Capture-C probes

To identify genes that are developmentally regulated during the fetal-to-adult transition, we analyzed RNA-Seq data of E14.5 fetal liver HSCs and adult bone marrow HSCs. Using EBSeq(*55*), we identified 7174 differentially expressed transcripts (corresponding to 3464 genes) at a false discovery rate (FDR) cutoff of 0.05. To focus on genes that are specifically expressed in HSCs, we compared the HSC RNA-Seq data to an RNA-Seq compendium of 100 mouse tissues/cells generated by the mouse ENCODE project.

Using a Z-score cutoff of 2, we identified 1921 transcripts that are highly expressed in both fetal liver and bone marrow HSCs. By overlapping the two sets of transcripts, we identified 715 transcripts that are expressed at high levels in HSCs and are differentially expressed between fetal liver and bone marrow HSCs. We also included additional 238 genes that are implicated in HSC biology based on literatures evidence(*56-61*). We merged transcripts whose TSSs are located in the same DpnII restriction fragment. In total, 4052 transcripts were selected (Supplemental Table S2). For each transcript, we defined the 1kb upstream and 1kb downstream of the TSS as the promoter region.

Capture probes for selected promoter regions were designed using the online tools (http://apps.molbiol.ox.ac.uk/CaptureC/cgi-bin/CapSequm.cgi)(*23,* 62). Briefly, the genomic coordinates of DpnII sites overlapping the target promoters were identified and 120-bp sequences from both Dpn II sites were generated for each fragment. Candidate probe sequences were filtered based on repeat density score <= 10 and simple repeat content <= 30. The remaining probe sequences were submitted to custom design of SureDesign capture oligos by Agilent.

### Capture-C

Capture-C assay was performed based on previous publication with a few modifications(*23, 62*). Half million FACS-sorted cells were cross-linked with 2% formaldehyde for 10 min at RT, quenched with glycine. The cross-linked cells were washed with pre-chilled PBS and lysed with 1mL cold lysis buffer (10 mM Tris, 10 mM NaCl, 0.5% NP-40, and 1x Protease inhibitors) for 10 min on ice. Nuclei were centrifuged at 1600x g for 5 min at 4°C and washed with ddH_2_O. The nuclei pellet was digested with three aliquots of 500 U DpnII and incubated at 37°C for 16∼24hr. DpnII was heat-inactivated by incubating samples at 65°C for 20 min. Chromatin fragments were ligated with 100 U T4 DNA ligase at 16°C for 8 hrs with slow rotation. Samples were de-cross-linked with 3 μL Proteinase K at 65°C overnight, followed by RNase A treatment for 30 min at 37°C. DNA was purified using phenol-chloroform extraction and precipitated using 70% ethanol.

3C DNA was sonicated to 200∼300bp using Covaris S220 ultra-sonicator (6 min; duty cycle, 10%; intensity, 5; cycle per burst, 200). Sequencing libraries were constructed using NEBNext Ultra Kit. The libraries were size-selected using AMPure XP beads.

Oligonucleotide capture was performed using the SureSelect XT2 protocol (G9621A, Agilent). The post-captured library was amplified with Herculase II Master Mix and purified using AMPure XP beads. The libraries were sequenced with 150 bp paired-end reads on Illumina NextSeq 500 or HiSeq 2000.

### ATAC-Seq and histone modification ChIP-Seq

Assay for Transposase-Accessible Chromatin using Sequencing (ATAC-Seq) was performed based on previous study with minor modification(*63*). Briefly, 50,000 FACS-sorted cells were centrifuged at 1600g for 5 min at 4°C, followed by one wash using 50 μL of pre-chilled 1x PBS and centrifugation at 1600g for 5 min at 4°C. Cells were lysed using pre-chilled lysis buffer (10 mM Tris-HCl, pH 7.4, 10mM NaCl, 3mM MgCl_2_ and 0.1% IGEPAL CA-630). Nuclei were centrifuged at 1600g for 10 min at 4°C. Nuclei were re-suspended in transposase reaction mix (25 μL 2×TD buffer, 2.5 μL transposase (FC-121-1030, Illumina) and 22.5 μL nuclease-free water) and incubated for 30 min at 37°C. The sample was purified using a Qiagen MinElute kit (28004, Qiagen). Following purification, libraries were amplified using 1x NEBNext PCR master mix and custom Nextera PCR primers. Libraries were size-selected at 100-700 bp by gel extraction (28604, Qiagen). Libraries were quantified with KAPA qPCR and bioanalyzer prior to pair-end sequencing on Illumina HiSeq 2000.

Low-Cell-Number ChIP-Seq was performed as following. Briefly, 50,000 cells for each IP were cross-linked with 1% formaldehyde (28906, Thermo Scientific) for 5 min at RT. Cells were re-suspended in 1x shearing buffer and sonicated with Covaris E220 for 780 seconds. 5% sheared chromatin was used as the input and the remaining chromatin was used for IP. IP was performed using ChIP-IT high sensitivity kit (53040, Active Motif) with some modifications. IP and input samples were treated with RNase A followed by proteinase K treatment. Cross-linking was reversed by incubating overnight at 65°C. Reverse crosslinked DNA was purified using a MinElute PCR purification kit (Qiagen, 28004) and re-suspended in 10 μL nuclease-free water. All IPed DNA and 1 ng input DNA were used for library preparation using the ThruPLEX-FDPrep kit (R40048, Rubicon Genomics) with 12 cycles of ampli?cation for IP DNA and 9 cycles for input DNA. Libraries were quantified with KAPA qPCR and bioanalyzer prior to single-end sequencing on Illumina HiSeq 2000.

### DNA fluorescence in situ hybridization (DNA-FISH)

Sorted cells were washed once with 1 mL PBS. Cells were fixed with 1 mL of MAA (methanol: acetic acid = 3:1) for 15 min on ice and spun down and re-suspended in 1 mL of MAA. This process is repeated for at least three times. Five million fixed cells were re-suspended in 1 mL of MAA. Cells were immobilized on the slide and denatured at 72°C for 3 min. Hybridization was performed in a dark humidity chamber for 3 days. Slides were washed with SSC buffer (S6639, Sigma), followed by staining with DAPI (P36935, Invitrogen). Slides were stored at -20°C or imaged immediately. Probes used in DNA-FISH experiments are listed in Supplemental Table S6.

### Cell culture

Hematopoietic precursor cell-7 (HPC-7) cells were grown in IMDM medium (12440-053, Invitrogen) with 10% FBS, 10% stem cell factor conditional medium, 1% Pen/Strep, and 7.48× 10^−5^ MTG (M6145, Sigma). Stem cell factor conditional medium was produced by BHK/NKL cell line. HPC-7 cells were maintained at the density of 5× 10^5^ ∼ 2 × 10^6^ cells per mL. OP9 and OP9-DL1 cells were grown in α-MEM (12571-063, Invitrogen) medium with 20% FBS, and 1% Pen/Strep.

### CRISPR-mediated knockout of *Tcf3* in HPC-7 cells

Guide RNA sequences targeting *Tcf3* were designed using Deskgen tools (Supplemental Table S6). Annealed sgRNAs were cloned into lentiCRISPR v2 (52961, Addgene). Lentivirus was produced by co-transfecting with pMD2.G (12259, Addgene) and psPAX2 (12260, Addgene) into HEK293FT cells. HPC-7 cells were transduced by lentivirus and positive cells were selected by culturing with 0.5 ug/mL puromycin for 21 days. Knockout of *Tcf3* was confirmed using Western Blot.

### Hi-C data processing

Hi-C read mapping, detection of valid interactions, correction of systematic noise, and calculation of normalized contact matrices were performed using HiC-Pro(*64*) with default parameters. Paired-end reads were mapped to the mm9 version of the mouse genome. Normalized contact matrices at 10kb resolution were computed using the ICE algorithm with default parameters.

### Analysis of chromatin compartments

Chromosome compartments were identified using principal component analysis (PCA). We first calculated the contact matrix for each chromosome using TAD as the unit. A cell of the contact matrix *O*_*ij*_ represents the total number of contacts between the *i*th and *j*th TADs. We adjusted the contact matrix according to the TAD sizes and distance as *O*_*ij*_/*s*_*i*_*s*_*j*_*E*_*ij*_, where *s*_*i*_, *s*_*j*_ and *E*_*ij*_ are the sizes of the *i*th *and j*th TADs, and the averaged contact frequency between genomic loci with distance *d*_*ij*_, which is the distance between the middle points of the *i*th and *j*th TADs. Next, we converted the above contact matrix to Pearson’s correlation matrices and PCA was conducted on the correlation matrices.

The sign of first principle component, denoted as PC1, was used to assign compartment label. Because the sign of PC1 was arbitrary, additional information was used for compartment assignment. As suggested by(*12, 66*), genomic regions with high gene density were assigned to positive PC1 values and correspond to compartment A. The rest of the genome were assigned to negative PC1 value and correspond to compartment B. Degree of compartmentalization was measured by the contact frequency between all possible pairs of TADs from the same (AA or BB) or from different types of compartments (AB). The normalized contact frequency of each pair of TADs was computed as the log2 ratio of the total number of observed inter-TAD contacts to the total number of expected inter-TAD interactions. The expected number of contacts between any pair of loci was calculated using the Shaman R package (https://bitbucket.org/tanaylab/shaman).

### Analysis of topologically associating domains (TADs)

TADs were called using normalized Hi-C data and the GMAP algorithm(*25*). Two TAD boundaries were considered shared if they are within 50kb of each other.

### Calculation of TAD boundary strength

To compute the score for TAD boundary strength, we first calculated the log2 ratio of observed to expected contact frequency between any two genome loci using the Shaman R package which we referred to as the Shaman ratio hereafter. The boundary strength score was then defined as the difference between the intra-TAD Shaman ratio and the inter-TAD Shaman ratio between the 600kb up-and down-stream regions flanking a TAD boundary. To identify boundaries with significantly altered strength between FL HSCs and BM HSCs, we first computed a null distribution of boundary strength difference using 40,000 randomly selected genomic loci that do not overlap with any observed TAD boundaries. The p-value for altered boundary strength was then computed based on the null distribution. Multiple testing correction was conducted using the Benjamini-Hochberg procedure. Dscore, a statistic provided by the Shaman package, was used to visualize TADs and TAD boundaries.

### Dynamics of promoter-centered intra-TAD interactions

We studied the dynamics of promoter-centered intra-TAD interactions by taking advantage of our high-resolution Capture-C data. Promoter-centered chromatin interactions were identified using the LiMACC algorithm with an FDR cutoff of 0.01. For each significant interaction identified in at least one cell type, the normalized interaction frequencies in both cell types were paired. TADs with fewer than 5 interaction pairs were excluded. P-values for TADs with significant changes in promoter interactions was computed using paired t-test. P-values were adjusted for multiple testing using the Benjamini-Hochberg procedure.

### 3D genome structure modelling

ICE (*65*) normalized Hi-C contact matrix was further normalized by quantile normalization between FL HSCs and BM HSCs at 25kb resolution. Whole chromosome models were reconstructed on the quantile normalized Hi-C contact matrix using LorDG(*67*). The method is robust to noise and inconsistency in Hi-C data. It works by first translating contact frequencies into spatial distances and then solving an optimization problem to build 3D models consistent with the spatial distances. The 3D coordinates of each binned locus were used for calculating Euclidean distance between two adjacent TADs associated with a given TAD boundary (Fig. 2F).

### Capture-C data processing

Capture-C data were processed using the pipeline from Hughes et al (*23*). Briefly, Illumina TruSeq adaptor sequences were trimmed from raw reads using trim galore version 0.41 with default parameter setting. Paired-end reads were merged into one single fastq file to ensure each pair of reads interleaved in strict order. Reads were mapped to the mouse genome (mm9) using Bowtie2 (v2.2.2). Mapped reads were analyzed using the script CCanalyser2.pl (https://github.com/telenius/captureC/releases). Unique informative reads were extracted for each captured bait as the input for calling significant interactions.

### Enhancer and promoter prediction using CSI-ANN

Enhancers and promoters were predicted using the CSI-ANN algorithm. The inputs to the algorithm are normalized ChIP-Seq signals of four histone marks (H3K4me1, H3K4me3, H3K27ac, and H3K27me3). The algorithm combines signals of all histone marks and uses an artificial neural network-based classifier to make predictions.

### Interaction calling using Capture-C data

We developed a **l**ocal **i**terative **m**odeling **a**pproach for identifying chromatin interactions using **C**apture-**C** or Capture Hi-C data (LiMACC). The basic idea is to categorize all capture bait and other end interactions (BOEIs) into different groups based on the distance between the two ends, and then fit a negative binomial model in each group. To estimate the null distribution, an iterative model fitting approach is used.

Suppose there are *m* baits, and *N* BOEIs, and each BOEI is supported by at least one read.

1. For each BOEI, calculate the distance between the corresponding bait and other end, as *d*_1_, *…*, *d*_*N*_.
2. Classify each BOEI into one of the following *G* groups based on the distance:
  a. Let *q*_*k*_ be the 100*k %* quantile of the *N* distances, *k = 0, 1,…G*, and
  b. The *i*th BOEI is pooled into group *k* if *q*_*k*–1_ ≤ *d*_*i*_ < *q*_*k*_.
3. To generate the null distribution, in each group, we iteratively define high confidence random contacts (HCRC) and fit a negative binomial distribution using the HCRCs.
4. Calculate raw p-values for each BOEI using the null distribution obtained from step 3 and pool the raw *N* p-values and adjust them based on the Independent Hypothesis Weighting (IHW) procedure (*68*).

The detailed iterative procedure for model fitting is given as follows:

1. In each group, define the initial HCRCs as those with the number of reads in the bottom 95 percentile.
2. Fit the null distribution in each group using the corresponding HCRCs and calculate the raw p-values.
3. Pool all raw p-values and adjust them by either the BH procedure or IHW procedure.
4. For each group, define HCRCs as those whose adjusted p-values are greater than a given FDR cutoff.
5. Repeat step 2-4 until there is no change in the set of HCRCs for each group and output the corresponding adjusted p-values for each interaction.

#### Normalization of raw read counts

Some BOEIs may have larger number of reads than others due to experimental biases. We therefore normalized the average number of reads per bait to a fixed number.

Suppose *N*_*ij*_ is the raw read count between bait *i* and the other end *j,* let 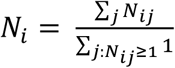 and *M* as the median number of {*N*_*k*_}_1≤*k* ≤*m*_, then the normalized reads are defined as:

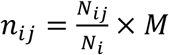

#### Adjusting bait to bait bias

Given the nature of the Capture-C protocols, the interactions between two baits are more likely to be captured than interactions between a bait and a non-bait. We propose to adjust this bait to bait bias in the following way:

For a given bait *i*, normalize the median contacts of *i* to another bait to be the same as the median contacts of bait *i* to other non-bait ends. We adjust such effect separately for each bait. Let *O*_*i*_ be the median number of {*n*_*ik*_} _*k is not a bait*_, and *B*_*i*_ be the median number of {*n*_*ik*_} _*k is also a bait*_ then we adjust *n*_*ij*_ between bait i and j as:

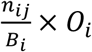

#### Promoter interacting regions (PIRs)

We re-binned the other end fragments of significant interactions into 2kb windows. The score of each window was defined as the largest LiMACC score (negative log-transformation in base 10 of the adjusted p-value) of all other ends that mapped to that window. We denoted those other end windows as Promoter Interacting Regions (PIRs). Downstream analyses such as transcription factor enrichment analysis and clustering analysis were based on PIRs.

### ChIP-Seq data processing

Sequencing reads were mapped to the mouse genome (mm9) using Bowtie2 (v2.2.2)(*69*) with default parameter setting. Uniquely mapped reads from both ChIPed and input DNA were used to compute a normalized signal for each 200 bp bin across the genome. Normalized signal is defined as following:

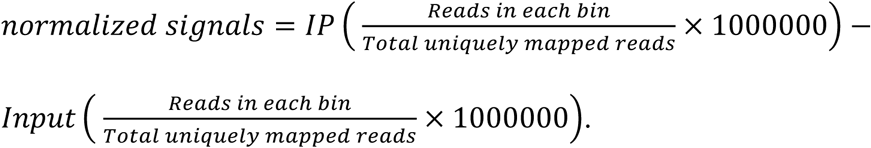

### ATAC-Seq data processing

Ilumina Nextera transposase adaptor sequences were trimmed from raw reads using trim galore version 0.41 with default parameter setting. Trimmed reads were mapped to the mouse genome (mm9) using Bowtie2 (v2.2.2) (*69*) and default parameter setting. ATAC-Seq peaks were called by MACS using default parameter setting.

### TF motif analysis of enhancers involved in cell-specific enhancer-promoter interaction

The DNA binding motifs of 718 TFs were downloaded from the CIS-BP database. The FIMO software(*70*) was used to scan the enhancer regions that overlap with ATAC-seq peaks. Significant motif hits were called using a p-value cutoff 0.01 with Bonferroni correction. Hypergeometric test was used to determine the enrichment of a given TF motif in the set of enhancers involved in stage-specific enhancer-promoter interactions. Raw hypergeometric p-values are corrected for multiple testing using the Benjamini-Hochberg procedure.

### Gene ontology analysis

GO term enrichment analyses were performed using the DAVID tool (*71*) (version 6.8). Raw p-values were adjusted using the Benjamin-Hochberg procedure.

### Availability of data and materials

All data generated in this study has been deposited in the Gene Expression Omnibus (GEO) database under the accession number GSE119201. LiMACC algorithm is freely available on GitHub (https://github.com/wbaopaul/limacc).

## Acknowledgments

We thank the Research Information Services at the Children’s Hospital of Philadelphia for providing computing support. This work was supported by National Institutes of Health of United States of America grants GM104369, GM108716, HG006130 (to K.T.), HD089245 (to K.T. and N.S.) and HL129998 (to G.B.).

## Author Contributions

Contribution: C.C, W.Y., and K.T. conceived and designed the study. C.C, H.Y., J.T., and P.G. performed research. W.Y. analyzed data. B.H., T.T., N.S., and B.G. provided additional experimental and analytical tools. K.T. supervised the study. C.C., W.Y., and K.T. wrote the paper.

